# Maize mutants in miR394-regulated genes show improved drought tolerance

**DOI:** 10.1101/2023.12.05.570184

**Authors:** Franco Miskevish, Anabella Lodeyro, María Agustina Ponso, Carlos Bouzo, Robert Meeley, Marja C. Timmermans, Marcela Dotto

**Author notes:** Corresponding author: Marcela C. Dotto Instituto de Ciencias Agropecuarias del Litoral (ICIAGRO-Litoral) Kreder 2805, Esperanza, Santa Fe, Argentina. CP3080 TE. Instituto Multidisciplinario de Investigación y Transferencia Agroalimentaria y Biotecnológica (IMITAB, CONICET-UNVM), Villa María, Córdoba, Argentina. **One sentence summary** Maize mutants in miR394-regulated genes show improved drought tolerance, due to more efficient intrinsic water use, changes in root architecture and increased epicuticular wax in drought conditions.

## Abstract

Water limitation represents one of the major threats to agricultural production, which often leads to drought stress and results in compromised growth, development and yield of crop species. Drought tolerance has been intensively studied in search for potential targets for molecular approaches to crop improvement. However, drought adaptive traits are complex and our understanding of the physiological and genetic basis of drought tolerance is still incomplete. The miR394-LCR pathway is a conserved regulatory module shown to participate in several aspects of plant growth and development, including stress response. Here we characterized the miR394 pathway in maize, which harbors two genetic loci producing an evolutionarily conserved mature zma-miR394 targeting two transcripts coding for F-Box proteins, named hereby ZmLCR1 and ZmLCR2. Arabidopsis plants overexpressing zma-MIR394B gene showed high tolerance to drought conditions, compared to control plants. Moreover, analysis of growth and development of single and double maize mutant plants in ZmLCR genes indicate that these mutations do not affect plant fitness when they grow in normal watering conditions, but mutants showed better survival than wild type plants under water deprivation conditions. This increased drought tolerance is based on a more efficient intrinsic water use, changes in root architecture and increased epicuticular wax content under water limiting conditions. Our results indicate that the miR394-regulated ZmLCR genes are involved in drought stress tolerance and are remarkable candidates for maize crop improvement.

## Introduction

One of the major threats to agricultural production is water limitation, affecting growth, development and yield of crop species world-wide. Water deficit impacts on several aspects of plant growth, affecting diverse functions such as transpiration, photosynthesis, leaf and root growth, as well as reproductive development (Nuccio *et al*., 2018; Tardieu, Simonneau and Muller, 2018; Martignago *et al*., 2020). Water saving in crop production under climate and environmental changing conditions represents a critical issue for agricultural economy and sustainability. In this regard, drought tolerance is one of the main targets for molecular approaches to crop improvement. Being produced in spring and summer, the maize growth cycle is characterized by a high-water demand (Gullì *et al*., 2015), therefore the development of varieties with higher stress tolerance and a lower water demand is highly desirable. However, drought adaptive traits are complex and multigenic and our understanding of the physiological and genetic basis of drought tolerance is still incomplete; resulting in rare availability of specific genetic targets (Roy, Tucker and Tester, 2011; Ruggiero *et al*., 2017; Manna *et al*., 2021).

Conservation across species of the miR394 pathway has been bioinformatically analyzed and found to be conserved in angiosperms (Kumar *et al*., 2019). This particular miRNA is 20-nt-long and is responsible for the post-transcriptional down-regulation of *LEAF CURLING RESPONSIVENESS (LCR)*, a gene encoding a F-Box protein, one of the components of E3 ubiquitin ligase complexes (Song et al., 2012; Knauer et al., 2013). However, a functional characterization of the components of this pathway is still missing in major agricultural crops such as maize, a species of high agronomical interest and value to world-wide crop production.

Studies performed mostly in Arabidopsis indicate that the miR394-LCR regulatory module is involved in a wide range of biological processes contributing to plant development, including stem cell competence in embryo shoot apical meristem (Knauer *et al*., 2013), regulation of flowering time (Bernardi *et al*., 2022), leaf morphology in Arabidopsis (Jian Bo Song *et al*., 2012), leaf inclination in rice (Qu, Li-Bi and Hong-Wei, 2019) and fruit and seed development in *Brassica napus* (Song *et al*., 2015). In addition, this pathway was shown to play roles in the response to biotic stress in tomato plants infected with *Botrytis cinerea* and *Phytophthora infestans* (Tian *et al*., 2018; Zhang *et al*., 2021), as well as in garlic plants infected with *Fusarium oxysporum* (Chand, Nanda and Joshi, 2016). Finally, a role in the response to abiotic stress has also been reported for the miR394-LCR module, with participation in drought and cold stress response in Arabidopsis (Song *et al*., 2013, 2016), while overexpression of miR394 from soybean (*Glycyne max*) and foxtail millet (*Setaria italica)* in Arabidopsis results in drought tolerant plants (Ni *et al*., 2012; Geng *et al*., 2021).

Considering that current knowledge about the importance of this regulatory pathway in agronomically important crops is limited, in the present work we characterized the miR394 pathway in maize. We identified two genetic loci in the maize genome producing a conserved mature zma-miR394, which regulate two target transcripts coding for F-Box proteins, named hereby ZmLCR1 and ZmLCR2. Analysis of growth and development of single and double *zmlcr* mutant plants indicate these mutations do not affect plant fitness negatively when they grow in normal watering conditions, but mutants showed better survival than wild type (WT) plants when grown in drought conditions. This increased drought tolerance is based on a higher photosynthetic activity in the mutants, resulting in a more efficient intrinsic water use (iWUE), changes in root architecture and increased epicuticular wax content under water limiting conditions. Our results indicate that the miR394-regulated ZmLCR genes are involved in drought tolerance in maize and the ZmLCR are remarkable gene target candidates for maize crop improvement.

## Results

### The components of the miR394 pathway in the maize genome

Previous work identified two MIR394 precursor genes in the maize genome (Zhang *et al*., 2009), which are annotated as such in version 3 of B73 reference genome (B73 RefGen v3). Here we updated genomic coordinates to the current version of the maize genome, summarized in Table S1. Both MIR394 transcripts present complex hairpin structures and give rise to the same conserved 20-nt mature miRNA (Fig. S1A). In order to identify the targets of miR394 in maize, we used TargetFinder (Allen et al., 2005) to predict genome-wide miR394 target genes and used available degradome data from the Next-Gen Sequence DBs (https://mpss.danforthcenter.org/dbs/index.php?SITE=maize_pare) to validate miR394 target transcripts in maize (Fig. 1A-B). We identified only two genes as putative miR394 targets, identified as Zm00001eb143020 and Zm00001eb364210 in the last maize genome annotation (Table S1), which have binding sites for miR394 (Fig. 1A). These genes code for F-Box family proteins and were named *ZmLCR1* and *ZmLCR2* in accordance to their miR394-regulated counterpart in Arabidopsis, *LCR* (Song et al., 2012). These two transcripts showed degradome signatures significantly above the baseline (Fig. 1B; arrows), corresponding to a cleavage event between nucleotides 10 and 11 in the perfectly complementary miR394 binding site (Fig. 1A, arrows).

**Figure 1:**
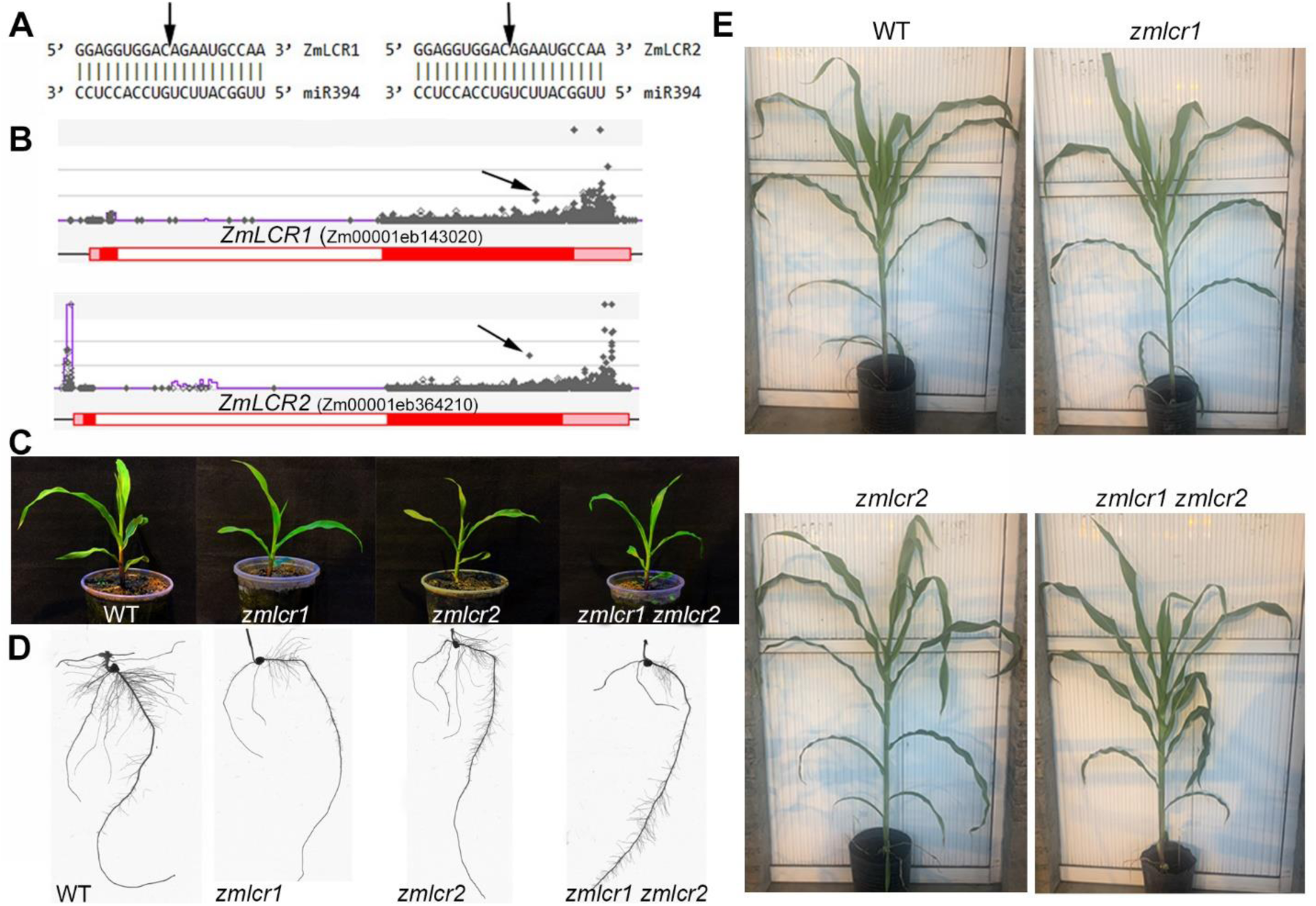
Target validation for maize miR394 and phenotypic analysis. A) Predicted binding site for miR394 on ZmLCR1 and ZmLCR2 transcripts. B) Degradome data visualization on ZmLCR1 and ZmLCR2 genomic region. Arrows in B) indicate reads supporting miR394 cleavage sites between the 10^th^ and 11^th^ nucleotide of miR394 on both target transcripts, indicated with vertical arrows in A). Graphics on B) were obtained from the Next-Gen Sequence DBs (https://mpss.danforthcenter.org/dbs/index.php?SITE=maize_pare). In both cases, normalized mapped reads are represented with grey rhombuses and basal accumulation of PARE signatures is indicated with a violet line. C) Seedlings of *zmlcr1*, *zmlcr2* and *zmlcr1 zmlcr2* mutants resemble WT plants. D) Roots from 10-days-old *zmlcr1*, *zmlcr2* and *zmlcr1 zmlcr2* mutants exhibit different root architecture from WT plants. E) Adult *zmlcr1*, *zmlcr2* and *zmlcr1 zmlcr2* mutant plants resemble WT plants, without differences in growth and development.

### Maize *zmlcr* mutants resemble wild-type plants phenotypically

To characterize the function of the miR394 pathway in maize, we analyzed growth and development of *zmlcr1* and *zmlcr2* single mutants, as well as *zmlcr1 zmlcr2* double mutant plants. The *zmlcr1* and *zmlcr2* are insertional mutants from the TUSC and UniformMu collections, respectively, which carry Mutator (Mu) transposon insertions (Meeley and Briggs, 1995; McCarty, Liu and Koch, 2018). The insertion is interrupting the coding sequence in the first exon of ZmLCR1, which was confirmed by sequencing (Fig. S1B), and is also the case for ZmLCR2 gene insertion (Fig. S1C). The location of these insertions allows for the assumption that these mutant lines are not able to produce a functional protein from these transcripts.

We compared WT and mutant plants throughout life cycle, but no developmental abnormalities were detected in the aerial part of the plants at the seedling stage when individuals of the three mutant genotypes exhibited leaves that resembled WT plants in terms of shape, width and did not show developmental abnormalities (Fig. 1C). Also, as they reached the adult stage, the mutant plants showed similar growth and development, being undistinguishable from their WT counterparts (Fig. 1E). Even though a different root morphology is observed in 10-days-old plants (Fig. 1D), showing mutant plants a decreased number of lateral roots, these differences are not maintained throughout growth and root architecture of adult mutant plants is similar to WT (Fig. 7, control conditions). These results indicate that ZmLCR genes might not be involved in leaf development and morphology in maize, which differs from Arabidopsis *lcr* mutants, shown to exhibit a leaf upward curling phenotype (Song et al. 2012).

### Maize miR394 pathway in the response to water-deprivation conditions

To further study the role of this pathway in maize, we analyzed miR394 and ZmLCR gene expression in maize plants exposed to drought conditions and observed a higher accumulation of this miRNA and the consequent lower expression of ZmLCR transcripts (Fig. 2A), indicating that the response to drought conditions in maize affects the expression of this regulatory module. In addition, and to evaluate the participation of zma-miR394 in the response to drought, we generated Arabidopsis transgenic plants overexpressing maize MIR394B gene (line named *OX-zma-MIR394B*), which accumulate 7.5 times more mature miR394 than control plants transformed with the empty vector (line named p402-EV), (Fig. S2A). We evaluated drought survival after 15 days without irrigation, followed by 7 days of re-watering, which resulted in the survival of 100% of the Arabidopsis transgenic plants in comparison to only 17% survival of control p402-EV plants (Fig. 2B). The behavior of Arabidopsis *OX-zmaMIR394B* plants resemble others overexpressing miR394 from soybean and foxtail millet (Ni *et al*., 2012; Geng *et al*., 2021), indicating the drought tolerant phenotype in Arabidopsis can be mediated by miR394 precursors from any of these three species. Interestingly, a general phenotypic evaluation showed *OX-zma-MIR394B* flower later than control p402-EV plants (Fig. S2B) and, even though no difference in primary root length was observed, a higher number of lateral roots was developed by *OX-zmaMIR394B* plants after 15 days of growth (Fig. S2C-D), indicating that this miRNA also participates in these others aspects of plant development.

**Figure 2:**
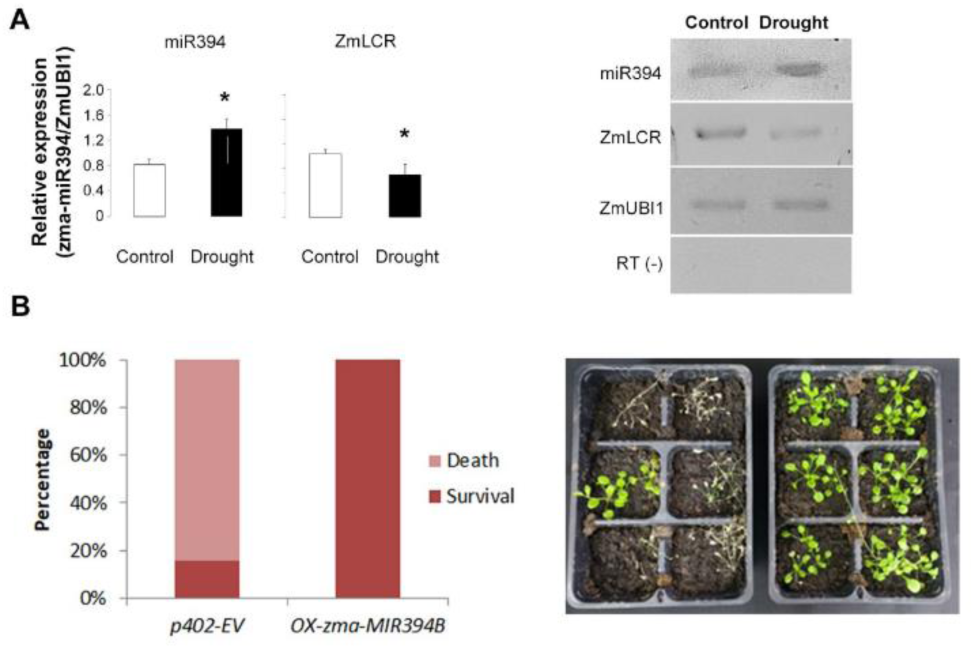
Maize miR394 in the response to drought conditions. A) miR394 accumulation and ZmLCR expression analyzed in WT plants grown in control, fully irrigated conditions or under drought conditions through water deprivation for 10 days. ZmUBI1: expression of maize UBIQUITIN 1 gene used as internal reference; miR394: expression of mature miR394 analyzed by Stem-loop RT-PCR; ZmLCR: expression of ZmLCR1 and ZmLCR2 genes using common primers to amplify both; RT (-): Retrotranscription negative control without enzyme. B) Percentage of survival and death for control p402-EV plants transformed with empty vector and *OX-zma-MIR394B* plants overexpressing maize Zma-MIR394B gene. Representative p402-EV and *OX-zma-MIR394B* plants are shown below, after 15 days of drought and 5 days of re-watering.

To further characterize the role of the miR394 pathway in the response to drought stress in maize, we performed a similar drought experiment for maize plants in which we analyzed the survival rate of WT and *zmlcr* mutant plants exposed to a long period of drought. Plants were grown under normal irrigation conditions until V3 stage and at this point watering was restrained for a period of 25 days. After 15 days without water, plants from all genotypes showed typical signs of water restriction, but the appearance of mutant plants of the three *zmlcr* mutant genotypes was noticeably better than WT plants (Fig. 3A). The latter showed leaf blades fully curled up, which is a typical response of maize plants to water limitation aiming to limit water loss through evaporation; whereas *zmlcr1*, *zmlcr2* and *zmlcr1 zmlcr2* plants still exhibited zones of expanded leaf blades (Fig. 3A, arrows). Water restriction continued for a total of 25 days, when all plants were similarly affected by the stress conditions and looked severely dry (Fig. 3B).

**Figure 3:**
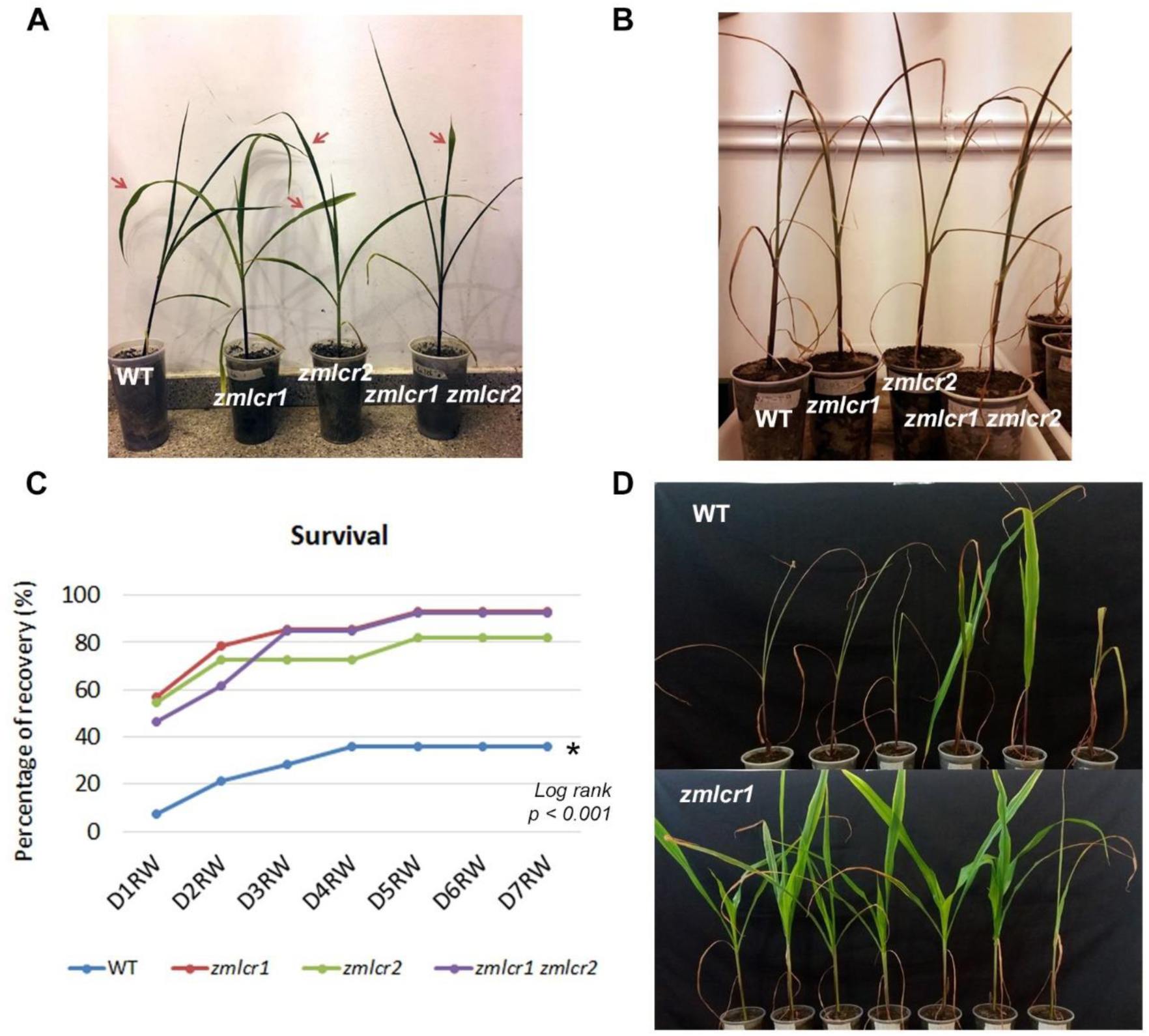
A) WT, *zmlcr1*, *zmlcr2* and *zmlcr1 zmlcr2* mutant plants after 15 days of water deprivation. WT plants exhibit fully curled up leaf blades, while mutant plants present regions of expanded leaf blades (arrows). B) WT, *zmlcr1*, *zmlcr2* and *zmlcr1 zmlcr2* mutant plants after 25 days of water deprivation. All plants look severely dry. C) Survival of WT, *zmlcr1*, *zmlcr2* and *zmlcr1 zmlcr2* mutant plants after re-watering during 7 days. Survival was assessed as the capability of plants to resume growth after re-watering. WT survival curve is different from the rest according to Kaplan-Meier log-rank analysis, followed by Holm-Sidak method for pairwise comparisons, P < 0.001. D) Representative WT and *zmlcr1* plants after 5 days of re-watering. Survivors re-open leaf blades, greenness return and plants resume growth.

Next, plants were re-watered during 7 days and plant survival was recorded as the percentage of plants from each genotype that were able to resume growth (Fig. 3C). While only about 36% of WT survived the imposed drought conditions, 92% of single *zmlcr1* and double *zmlcr1 zmlcr2* mutants recovered, as well as 82% of *zmlcr2* plants (Fig. 3C). Representative individuals corresponding to WT and *zmlcr1* genotypes are shown in Fig. 3D after 5 days of re-watering. These interesting results indicate that mutations in ZmLCR genes are beneficial for maize drought tolerance and survival to long drought periods and that the miR394 pathway participates in the response to drought conditions in maize.

### Mutations do not affect plant fitness in normal watering conditions but confer increased water use efficiency under drought

To further characterize the influence of these mutations on plant fitness, we compared plant height as an indication of plant growth for WT, *zmlcr1*, *zmlcr2* and *zmlcr1 zmlcr2* plants as well as physiological traits that allow for the study and comparison of the four analyzed genotypes, both under full irrigation conditions (Control) and under water deprivation (Drought), (Fig. 4). Interestingly, we determined there are no differences in any of the analyzed traits between WT and each of the mutant genotypes analyzed when plants are grown under normal watering conditions (Fig. 4, blue bars). Our analysis showed no difference in plant height (Fig. 4A, blue bars) or leaf transpiration (E, Fig. 4B, blue bars) and the greenness index values are also similar for all the genotypes (Fig. 4C, blue bars), indicating chlorophyll content is not affected by these mutations. Moreover, we detected no differences in stomatal conductance (gs, Fig. 4D, blue bars), net photosynthesis (Pn, Fig. 4E, blue bars) and intrinsic water use efficiency (iWUE = Pn/gs, Fig. 4F, blue bars) when compared to WT or among mutant genotypes. Taken together, the results presented so far indicate that these mutations do not affect plant development or growth significantly, allowing the plants to perform their basic physiological functions and maintain normal plant fitness when they grow in normal watering conditions.

**Figure 4:**
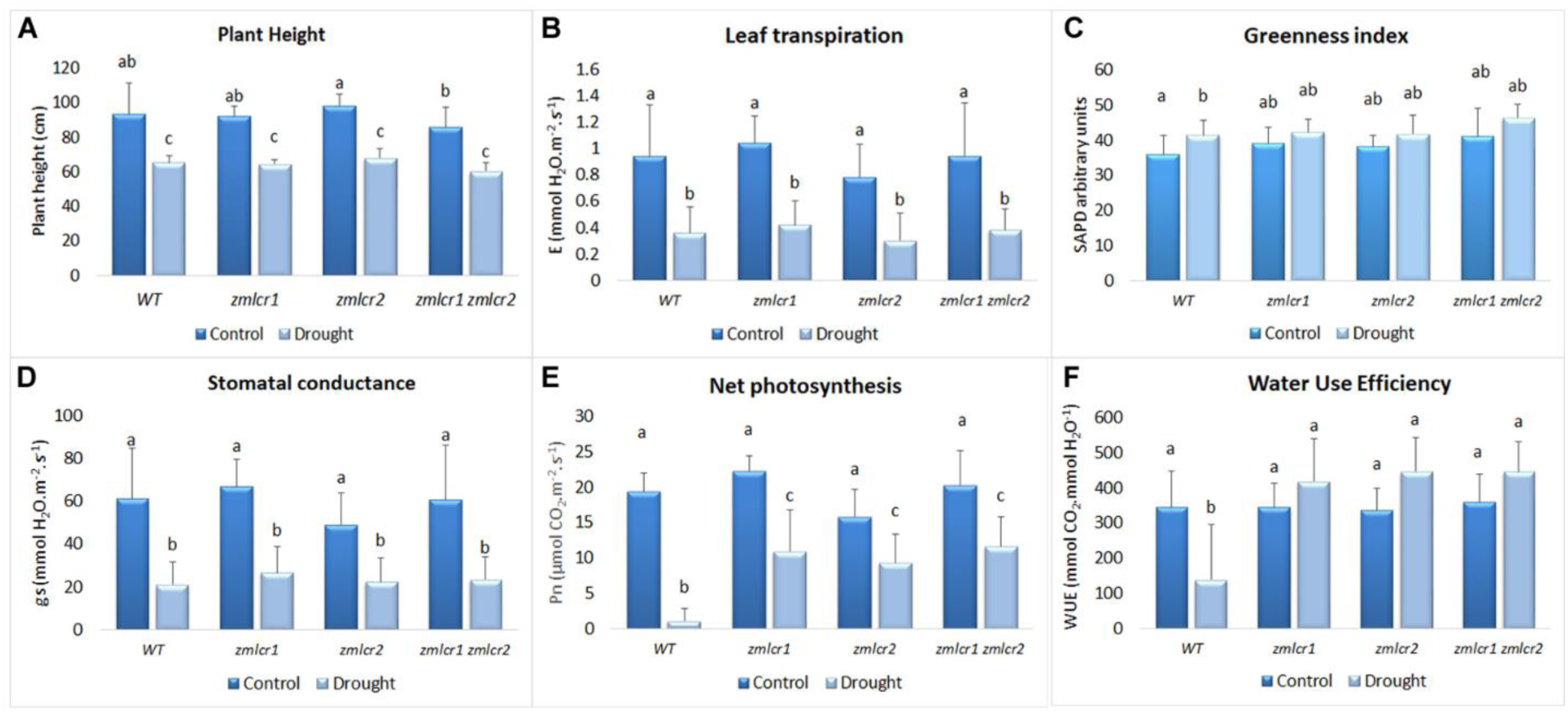
Growth and physiology of WT, *zmlcr1*, *zmlcr2* and *zmlcr1 zmlcr2* mutant plants in normal watering (Control) and after 15 days of water deprivation (Drought). A) Plant height measured from base of the stem to the tip of the longest leaf. B) Leaf transpiration (E, mmol H_2_O.m^-1^.s^-1^). C) Greenness index, measured using SPAD. D) Stomatal conductance (gs, mmol H_2_O.m^-1^.s^-1^). E) Net photosynthesis (Pn, μmol CO_2_.m^-1^.s^-1^). F) Intrinsic Water use efficiency (iWUE = Pn/gs x 1000, mmol CO_2_.mmol H_2_O^-1^). E, gs and Pn were measured simultaneously *in vivo* using a portable CIRAS-2 analyzer. The values are means ± SD (n ≥ 5). Different letters indicate significant difference using ANOVA followed by Tukey’s post-hoc comparisons when P ≤ 0.05.

Simultaneously, in order to understand the basis of the increased drought survival observed for mutant plants, we studied plant fitness by measuring and analyzing the same traits in plants without irrigation for 15 days (Fig. 4, light blue bars). We observed no difference among the four genotypes in plant height (Fig. 4A, light blue bars), leaf transpiration (E, Fig. 4B, light blue bars), greenness index (Fig. 4C, light blue bars), and stomatal conductance (gs, Fig. 4D, light blue bars), indicating that these mutations do not confer any advantage nor disadvantage for these traits under water limiting conditions. Interestingly, we determined net photosynthesis values are higher for the three mutants in comparison to WT plants (Pn, Fig. 4E, light blue bars) and when we calculated the intrinsic water use efficiency (iWUE, Fig. 4F, light blue bars), we observed that all three mutant genotypes use water more efficiently that WT plants (Fig. 4F, light blue bars). These results suggest that the increased drought survival observed for the *zmlcr* mutants, not observed in WT plants (Fig. 3C), is related to a more efficient water use based on increased net photosynthesis performance when plants are exposed to drought conditions.

In addition, most of the analyzed physiological parameters show, as expected, lower values in plants grown under water-limiting conditions compared to their counterparts with full irrigation (Fig. 4, control vs. drought): plants slow down growth under drought conditions, as evidenced by the lower height observed for all genotypes (Fig. 4A), leaf transpiration is also lower for all analyzed genotypes (Fig. 4B), as well as stomatal conductance (gs, Fig. 4D). However, SPAD-measured chlorophyll content is not significantly different from WT plants both under normal or water-limiting conditions and values remain comparable across both normal watering and drought conditions (Fig. 4C), indicating plants do not lose their green color at this point (Fig. 3A). Interestingly, mutant plants exhibited lower net photosynthesis values under drought conditions, but the decrease observed for WT plants was much more severe, reaching a level of Pn close to zero in drought-exposed plants (Pn, Fig. 4E). Particularly interesting is the result regarding iWUE values (Fig. 4F), which for mutant plants in drought conditions are similar to those observed in well-irrigated plants, contrary to WT plants which exhibit a lower iWUE in water-deprived conditions (Fig. 4F). These results suggest that mutations in ZmLCR genes improve water use efficiency by allowing plants to maintain a higher photosynthetic activity in spite of the absence of water, which negatively affects Pn and iWUE earlier in WT.

### Mutations in ZmLCR genes do not modify PSII photochemical capacity

To further analyze the photosynthetic performance of the mutant plants, chlorophyll fluorescence-based photosynthetic parameters associated with PSII photochemical capacity were measured. The different parameters were analyzed for WT, *zmlcr1, zmlcr2* and *zmlcr1 zmlcr2* double mutant plants in a 25-days-long drought experiment, measuring at different time points until plants appeared too dry to allow stable readings. No change in the quantum yield of photosystem II (PhiII) was observed in control conditions for any of the genotypes analyzed. However, this parameter declined steadily from the 19^th^ day of water deprivation in all lines (Fig. 5A). Even though values in PhiII decreased towards the end of the drought experiment, which is typically observed in plants under stress, there is no statistical difference between PhiII values obtained for WT and mutant plants (Fig. 5A). Since PhiII is a measure of the electron flow through photosystem II (PS II), a similar trend was observed for LEF (Linear Electron Flow) parameter, with lower values towards the end of the drought experiment for all genotypes under drought conditions (Fig. S3A). The maximum value of PhiII, represented by the Fv’/Fm’ ratio in light adapted leaves, reflects PSII integrity, and it was used to monitor permanent damage in PSII and their associated Light Harvesting complexes during the treatment (Baker, 2008). These values dropped from ∼0.7 to less than 0.6 in all lines at the end of the drought regime, indicating damage to PSII (Table S3) and severe drought stress level (Sommer *et al*., 2023).

**Figure 5:**
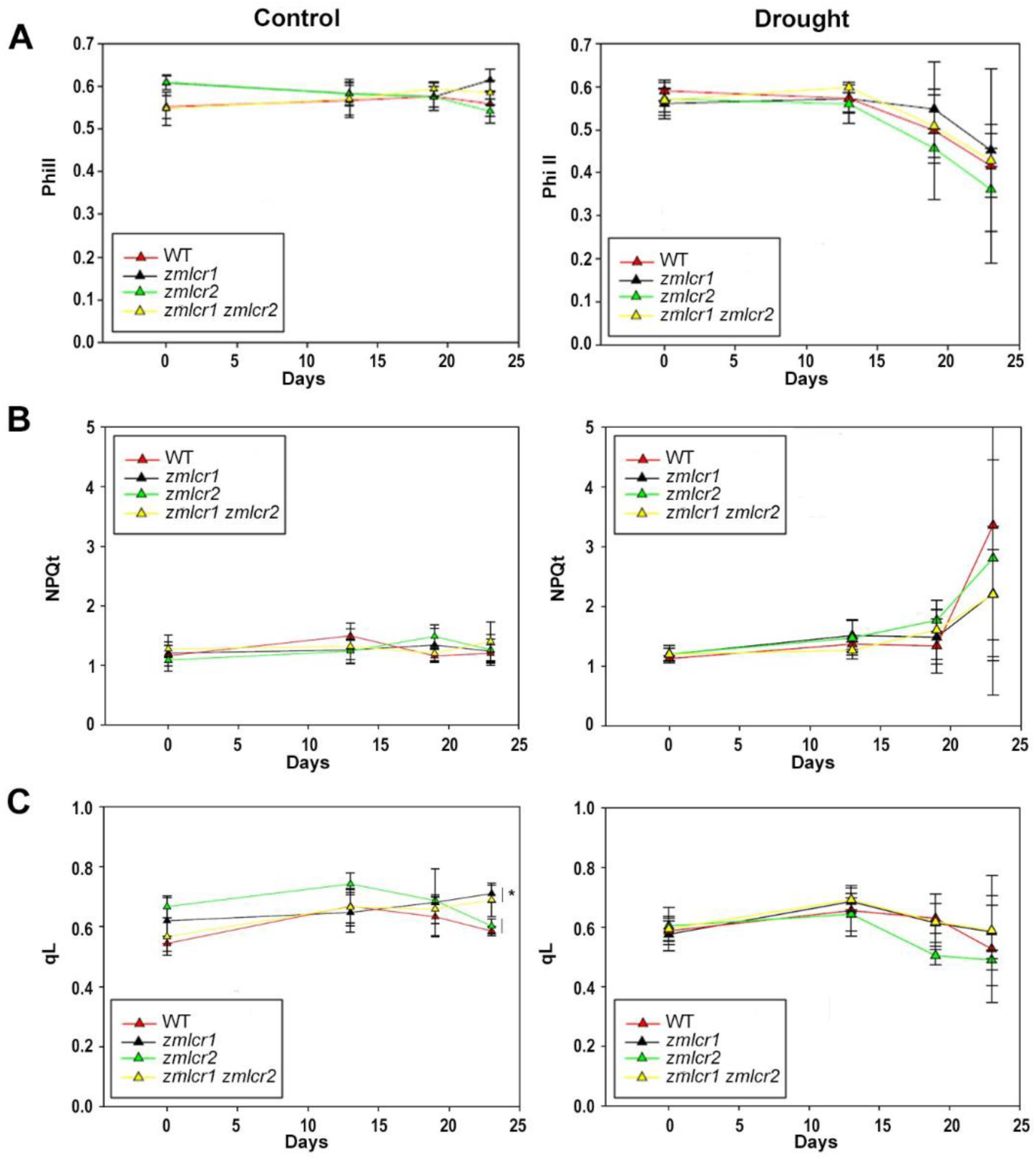
Energy management at PSII in WT, *zmlcr1, zmlcr2* and *zmlcr1 zmlcr2* mutants. Evolution of PhiII, NPQt and qL in fully-irrigated plants (Control) and water-deprived (Drought).

Water deprivation had effect on PS II energy dissipation processes, only at the end of the drought experiment (Fig. 5B), as it is reflected by the increased in the non-photochemical quenching (NPQt). Parameters NPQt and PhiNPQ estimate the extent of energy dissipation by heat (and hence the excess of excitation energy on the photosynthetic electron transport chain) when plants are stressed. No effect of the mutations was observed in comparison to the behavior of WT plants in both parameters (Fig. 5B and S3B, drought). A significant statistical difference was observed between WT and both *zmlcr1* and *zmlcr2* mutants at the beginning of the experiment for PhiNO parameter (Fig. S3C), which represents the fraction of light lost via non-regulated energy dissipation processes, suggesting that mutations could have an influence in lowering the constitutive loss of energy by the photosynthetic machinery. However, this difference was not observed at later points of the experiment, nor in control neither in drought conditions (Fig. ·S3C).

Finally, simple *zmlcr1* and double *zmlcr1 zmlcr2* mutant plants showed higher qL values than WT in the last measurement taken in control conditions (Fig. 5C). This parameter indicates the fraction of photochemically active centers in PSII, for which higher values could represent an indication of a more beneficial status for photosynthetic electron flow. However, this difference was not observed under drought conditions, being qL values approximately without changes for all genotypes along the experiment (Fig. 5C).

### Mutants in ZmLCR genes accumulate more epicuticular wax under drought conditions

In order to evaluate factors influencing water use and content in mutant plants, we analyzed relative water content (RWC), ion leakage and epicuticular wax content in well-watered and drought-treated WT and *zmlcr1 zmlcr2* double mutant plants (Fig. 6). Even though we determined both genotypes had a lower RWC under drought conditions, the difference between control and drought conditions was smaller for double mutant plants (Fig. 6A), suggesting mutations could help against water loss in plants growing with water restriction.

**Figure 6:**
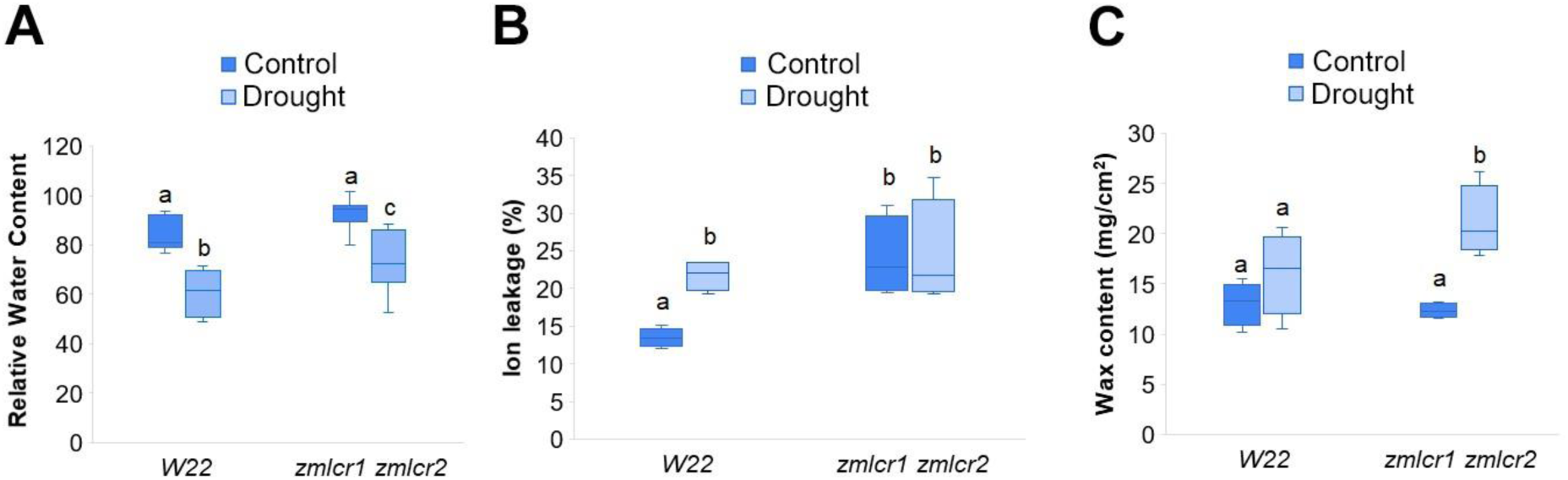
A) Relative water content, B) Ion leakage and C) Wax content determined for WT plants and *zmlcr1 zmlcr2* double mutants in control and drought conditions.

Next, we used ion leakage determination to further understand whether membrane integrity was affected differentially in WT and double mutant plants. We observed that *zmlcr1 zmlcr2* double mutant plants had a higher ion leakage percentage that WT plants in control conditions and that this value does not change when plants are grown under drought conditions (Fig. 6B). This result is intriguing since it suggests that mutations affect membrane integrity negatively, even in normal watering conditions, but this does not appear to affect plant development and physiology as shown in previous experiments presented here.

Finally, when we analyzed epicuticular wax content we observed no differential accumulation in WT plants under control or drought conditions and wax content was also similar in well-watered double mutant plants. Interestingly, in *zmlcr1 zmlcr2* double mutant plants we observed a higher wax accumulation for plants exposed to drought conditions (Fig. 6C), suggesting that this could contribute to a lower water loss in water-limiting conditions and possibly to a more efficient water use.

### Mutants in ZmLCR genes exhibit different root architecture under drought conditions

To analyze additional factors that could be contributing to the improved drought survival observed for *zmlcr* mutants (Fig. 3C), we analyzed root architecture in WT and double mutant plants, both under normal watering and drought conditions using WinRhizo scanner and software (Fig. 7). We did not observe differences in the general architecture between WT and *zmlcr1 zmlcr2* double mutants grown under control conditions (Fig. 7A, Control), whereas double mutant plants exhibited roots with increased formation of lateral roots (Fig. 7B, Drought). To quantify these observations, we analyzed ‘total root length’ as a general indication of root growth and ‘number of tips’ and ‘total length of lateral roots’ to compare secondary root growth (Fig. 7B-D). We determined that total root length does not change significantly in WT plants grown in control or drought conditions, but double mutant plants increase their total root length when exposed to drought conditions (Fig. 7B). This difference is also observed for *zmlcr1 zmlcr2* double mutant plants but not for WT plants when analyzing number of tips (Fig. 7C) and total length of lateral roots (Fig. 7D) under drought. This modification in root architecture observed for double mutant plants could also be influencing the higher drought tolerance after long periods of water restriction.

**Figure 7:**
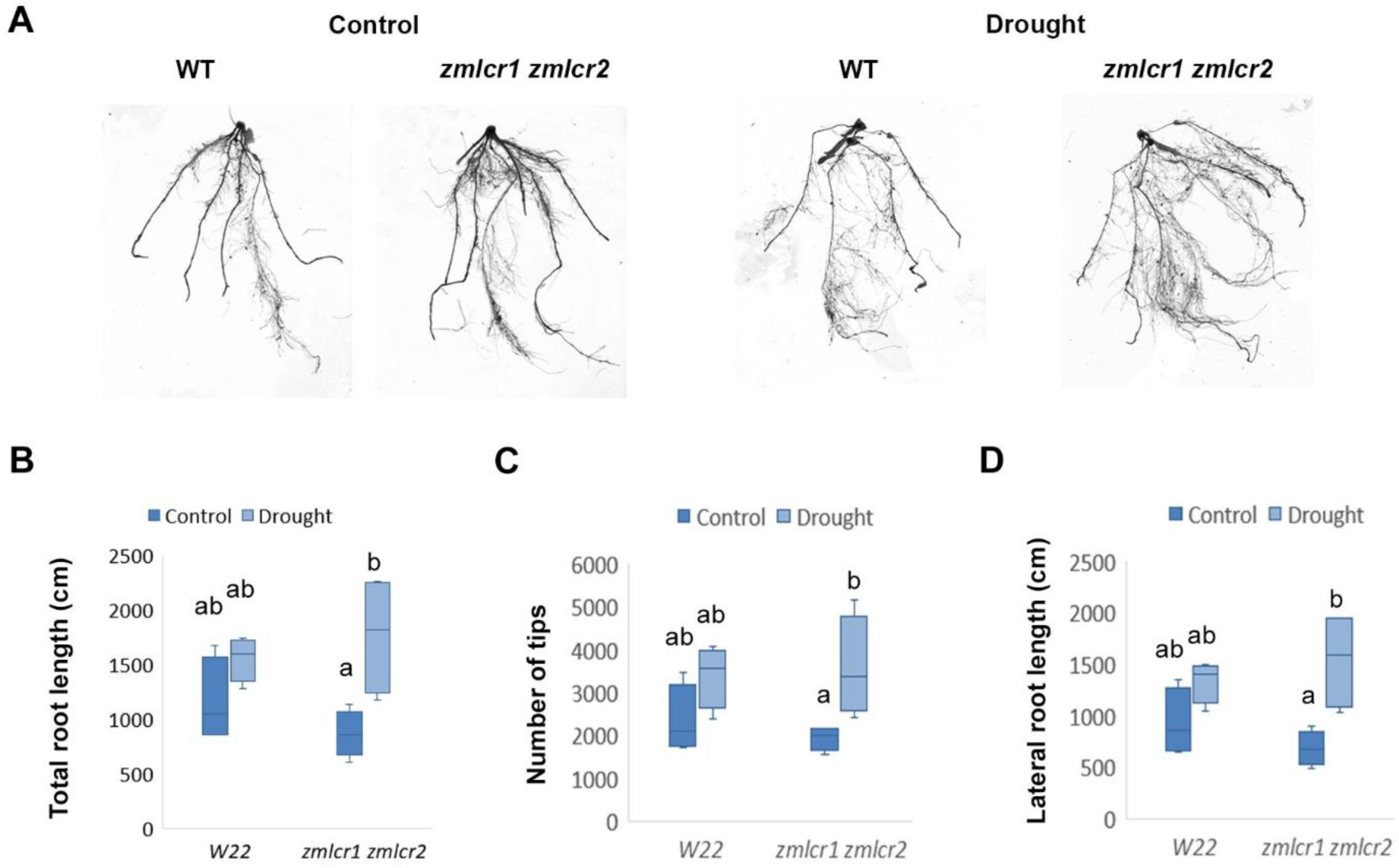
Root architecture analysis. A) Representative root systems from WT and *zmlcr1 zmlcr2* double mutant plants grown under normal irrigation (Control) and water-deprived conditions (Drought). B) Total root length C) Number of tips. D) Lateral root length. Analysis was performed using WinRhizo Scanner and Root Image Analysis System.

## Discussion

Genome-wide analysis of miRNA regulation mediated by transcript cleavage is possible through genome-wide target prediction combined with analysis of cleaved transcripts in degradome data obtained from sequencing and mapping of PARE libraries. This analysis detects above background accumulation of degradome reads that match predicted miRNA-mediated cleavage sites in possible target transcripts (Allen *et al*., 2005; German *et al*., 2008). In the present work we report the identification and characterization of two genes coding for F-Box proteins, regulated at the post-transcriptional level by miR394 in maize, as shown by the abundance of PARE signatures in degradome libraries (Fig. 1). Same as Arabidopsis, maize genome harbors two genetic loci coding for miR394 (Table S2), but in maize this miRNA is capable of regulating expression of two transcripts, ZmLCR1 and ZmLCR2, whereas a single AtLCR target gene has been reported in Arabidopsis (J. B. Song *et al*., 2012; Knauer *et al*., 2013). Gene duplication in plants is a widely-spread phenomenon, contributing in some cases to the evolution of novel molecular functions and providing functional redundancy in other examples (Panchy, Lehti-Shiu and Shiu, 2016; Li *et al*., 2021). Even though additional experiments are necessary to determine ZmLCR1 and ZmLCR2 molecular function, a role for ZmLCR1 and ZmLCR2 can be suggested as part of E3 ubiquitin-ligase complexes due to the presence of the conserved F-box domain in their predicted protein structure. E3 ubiquitin-ligases are SKP1-CULLIN1-FBOX (SCF) complexes responsible for ubiquitination of specific target proteins, where such specificity is determined by the F-Box component, resulting in proteins targeted for degradation by the 26S proteasome (Lechner *et al*., 2006; Chen and Hellmann, 2013; Shu and Yang, 2017). In Arabidopsis, the miR394/LCR module was shown to be important to define a domain of stem cell competence and for the development of proper leaf morphology (J. B. Song *et al*., 2012; Knauer *et al*., 2013). In maize, in situ hybridization showed adaxial expression of mi394 in maize leaf primordia, but no accumulation of miR394 in maize shoot apical meristem was detected (Ma *et al*., 2021). However, our characterization of *zmlcr* mutants did not reveal abnormalities in leaves (Figs. 1C, E) or any other developmental aspects. Instead, our results suggests that functional diversification of the miR394/LCR module exist between Arabidopsis and maize. For example, LCR-containing SCF complexes in Arabidopsis were shown to target members of the MAJOR LATEX PROTEIN family (MLP) for degradation (Litholdo *et al*., 2016). Whether this is also the case in maize remains to be determined, as well as whether ZmLCR1 and ZmLCR2 proteins are part of SCF complexes with the same or different target proteins. Our results showing there is no additive effect on drought tolerance for the simple and double *zmlcr* mutants are intriguing (Fig. 3C) and could be explained if ZmLCR-containing SCF complexes for each F-Box protein would target for degradation the same protein(s), which would accumulate to a functional higher level in either single or double mutants, enough to trigger enhanced drought tolerance. The specific molecular mechanism behind this response needs further work and a combination of biochemical and proteomic approaches.

We showed that miR394 expression is higher in maize plants grown under drought conditions, with the consequently decrease in ZmLCR gene expression (Fig. 2A), leading us to overexpress maize MIR394B gene in Arabidopsis, a simpler heterologous system that allowed us to explore the role of zma-miR394. Interestingly, Arabidopsis *OX-zma-MIR394B* plants showed a late flowering phenotype (Fig. S2B) which is in agreement with our previous publication demonstrating a role of miR394 in the regulation of flowering time in *A. thaliana* (Bernardi *et al*., 2022) and another work in which late flowering was also observed for *Brasssica napus* transgenic 35S::BnMIR394B plants (Song *et al*., 2015). Moreover, the study of *OX-zma-MIR394B* plants suggest a possible role for zma-miR394 in lateral root development, since a higher number was observed in 15-days-old transgenic plants (Fig. S2D). Regarding drought tolerance, *OX-zma-MIR394B* plants fully recovered after a 15-days-long drought period (Fig. 2B) which indicates zma-miR394 can be successfully processed from its precursor transcript in Arabidopsis plants as determined by mature miR394 expression analysis (Fig. S2A) and that a higher accumulation of this miRNA is capable of triggering a high drought resistance, similar to previous reports using Arabidopsis to overexpress miR394 from soybean and foxtail millet plants (Ni *et al*., 2012; Geng *et al*., 2021).

Best candidate genes for modification to improve agronomically important traits should cause minimum or null penalties when plants grow in favorable conditions, nor be associated with negative pleiotropic effects on other important agronomic or marketable attributes or other economically important crop qualities (Richards, 2006; Singh *et al*., 2020; Gao, 2021). A remarkable feature of the mutations in ZmLCR genes is that they do not affect plant development, therefore mutant plants have a normal appearance (Fig. 1C, E). Importantly, these mutations do not have a detrimental effect on plant fitness and physiology when grown in normal irrigation conditions (Fig. 4). These are highly desirable attributes for genetic crop improvement, since it guarantees beneficial changes in plants which will manifest under specific conditions, as observed for the higher survival rate in *zmlcr* mutants compared to WT after a long period of drought (Fig. 3C).

Our comprehensive analysis of the physiology of WT and mutant plants in normal and water-restricted conditions provide insights into several aspects of plant fitness. Differences in chlorophyll content are indicative of the strength of the internal leaf structure necessary for photosynthesis, which can be analyzed in a non-destructive way using SPAD chlorophyll meter (Markwell, Osterman and Mitchell, 1995). In our work we observed no differences between mutant and WT plants within neither growth condition and interestingly, there are no differences in chlorophyl content for mutant plants compared across conditions (Fig. 4C), indicating that drought conditions do not affect chlorophyll content negatively for *zmlcr* mutants in comparison to fully irrigated plants in our experimental conditions, but this is also the case for WT plants.

Our results show that *zmlcr* mutant plants exhibit a later leaf curling response in comparison to WT plants (Fig. 3A, arrows), which is a typical response to avoid water loss through evaporation when plants are grown in water-limiting conditions (Song, Jin and He, 2019; Wang *et al*., 2020). The later appearance of this response in mutant plants suggests that mutant plants are perceiving the drought condition at a later point, which could be related to the higher WUE observed for mutant plants when exposed to drought conditions (Fig. 4F). WUE as the ratio between Pn and gs is referred to as intrinsic WUE, which reflects net carbon assimilation per unit of water, representing a central aspect to analyze when trying to achieve higher WUE for crop improvement (Franks *et al*., 2015; Flexas *et al*., 2016; Hatfield and Dold, 2019). Typically, a higher WUE value results from lower stomatal conductance, but this is often linked to decreased photosynthetic capacity, reducing plant growth and productivity (Lawson and Blatt, 2014). Our results here show that *zmlcr* mutants are excellent candidates for further studies since we observed a lower stomatal conductance in drought conditions for all analyzed genotypes, as well as a lower net photosynthesis, but the decrease in Pn was smaller for *zmlcr* mutants than for WT plants (Fig. 4D-E), resulting in higher iWUE detected for *zmlcr* mutants in drought, which shows no difference with iWUE values measured in normal watering conditions (Fig. 4F). Altogether, these results indicate that these mutations make plants capable of using water in drought conditions as efficiently as in normal irrigation conditions, which is driven by a milder impact on photosynthetic activity compared to plants without these mutations.

In spite of the effect observed on Pn under drought conditions, our analysis of chlorophyll fluorescence-based parameters associated to PSII functionality did not show statistical differences between WT and any of the *zmlcr* mutants under drought conditions. Probably, as a consequence of a reduced metabolism and stomatal closure, drought triggered in both, wild type and mutant plants, a higher dissipation of absorbed light via NPQt at the end of the period of water deprivation (Figs. 5 and S3). A differential performance of the photosynthetic machinery could have explained the better survival exhibited by *zmlcr* mutants, since the analyzed parameters are tightly related to plant fitness and response to stress conditions (Murchie and Lawson, 2013; Gómez *et al*., 2018). However, the absence of differences between WT and *zmlcr* mutant plants indicate that damage in PSII and the photosynthetic electronic transport components due to the drought conditions occur at the same extent irrespective of the genotypes. Reduction in photosynthesis due to drought stress is generally attributed to destruction of photosynthetic pigments, changes in activity of PSII and PSI, reduction in stomatal conduction, influx of CO_2_ through stomata and its fixation by Calvin Cycle enzymes (Lawlor and Tezara, 2009). During the treatment, fluorescent measurements were used to follow the photoinhibitory effect of drought. The decrease of PhiII under drought stress has been reported, which is indicative of photo-inhibitory damage (Athar and Ashraf, 2005; Lawlor and Tezara, 2009), however the action of alternative sinks and metabolic pathways may, under certain conditions, affect the ability of chlorophyll *a* fluorescence to track the dynamics of photosynthetic CO_2_ assimilation (Porcar-Castell *et al*., 2014). Since the approach used here allows for specific events occurring at PSII, we cannot rule out a differential photochemistry at the level of PSI for WT and *zmlcr* mutants that could support the higher drought survival observed for the mutants (Suzuki *et al*., 2021).

Drought tolerance is generally related to enhanced capacity to maintain cellular and physiological functions when plants are growing in a water-limiting environment (Leakey *et al*., 2019). Plant’s response to drought is complex and different scenarios can occur that could explain the improved drought tolerance observed for these mutants (Fig. 3C). Physiological mechanisms that allow plants to maintain a high plant water status or improve drought tolerance by increasing dehydration avoidance are some of the possibilities (Morison *et al*., 2008; Bramley, Turner and Siddique, 2013). Here we showed that *zmlcr1 zmlcr2* double mutant plants showed a higher epicuticular wax content under drought conditions (Fig. 6C), which can help avoid dehydration and improve water availability for a more efficient use, contributing to the tolerant phenotype observed. In addition, our results indicate that the higher development of lateral roots can also contribute to the differential tolerance of mutant plants to drought (Fig. 7). Roots are usually the first organs to sense a water shortage, therefore changes in root architecture during drought periods typically occur in order to improve access to soil water. Common changes in root system include a deeper root system as well as an increase in lateral root development (Ribaut *et al*., 2009; Hazman and Kabil, 2022; Kou, Han and Kang, 2022)

As mentioned above, ZmLCR genes encode F-Box proteins that are part of SCF complexes targeting specific proteins for degradation by the proteasome. The definitive molecular mechanism behind the improved drought tolerance for maize *zmlcr* mutants requires the identification of these proteins which are likely to be key players in the response to drought. However, this is not an easy task since candidate target proteins are not available for maize at the moment. Despite this, our results identified *zmlcr* mutants as remarkable candidates for further exploration of agronomically important aspects related to plant growth and productivity in drought conditions. With null penalties in plant fitness under normal irrigation conditions and a higher iWUE when growing in drought conditions, these mutants are promising candidates for further studies to establish whether these mutations affect plant productivity and to evaluate their response to drought in field conditions.

## Materials and methods

### Plant material

Maize (*Zea mays* L.) mutant seeds in ZmLCR1 gene (GRMZM2G119650; Zm00001d042309) were obtained from Pioneer DuPont’s (Corteva’s) TUSC collection (TUSC BT94 177 B-03). Seed stock with insertions in ZmLCR2 gene (GRMZM2G064954; Zm00001d011994) were obtained from the UniformMu collection (seed stock ufmu-04186, locus mu1039927). Mutant plants were identified by PCR genotyping using gene specific primers (GSP) and GSP-MuTIR combinations (Table S1). Plants carrying each insertional mutation were introgressed three times into W22 background followed by self-pollination and identification of heterozygous and homozygous single mutants for each gene (*zmlcr1/zmlcr1* and *zmlcr2/zmlcr2*). Next, *zmlcr1/zmlcr1* mutants were crossed to *zmlcr2/+* plants, followed by identification of *zmlcr1/+ zmlcr2/+* individuals, which were next crossed to *zmlcr1/zmlcr1* mutants. The progeny was genotyped to identify *zmlcr1/zmlcr1 zmlcr2/+* individuals for self-pollination and identification of *zmlcr1/zmlcr1 zmlcr2/zmlcr2* double mutants, which were finally bulked. For simplicity, homozygous single and double mutant plants will be referred to as *zmlcr1*, *zmlcr2* and *zmlcr1 zmlcr2*.

### Arabidopsis transgenic plants generation

Vector pGWB402Ω (Nakagawa *et al*., 2007) was used to generate the *OX-zma-MIR394B* construct, which consists of 2X35S-CaMV promoter driving expression of zma-MIR394B gene. A total of 395 bp was amplified using specific primers (zma-394b-F/zma-mir394b-R; Table S1) and cloned into pCR8-GW vector (Thermo Fisher). The amplified region includes the annotated 122 bp hairpin (GRMZM5G871952) and additional upstream and downstream flanking sequence. Sanger sequencing was used to confirm the cloned sequence and positive clones were used in LR Gateway reaction into pGWB402Ω binary vector. Electrocompetent *Agrobacterium tumefaciens* cells, strain GV3101, were transformed and used to transform Col-0 plants using floral dip protocol (Clough and Bent, 1998).

### microRNA target prediction and validation

Target prediction for miR394 was made using Target Finder, allowing a maximum score = 4.5 (Allen *et al*., 2005) and using the full set of maize cDNA (Zea_mays.AGPv3.22.cdna.all) as possible targets. The PARE signatures corresponding to the predicted cleavage sites were analyzed using degradome data available at the Maize Next-Gen Sequence DBs (https://mpss.danforthcenter.org/dbs/index.php?SITE=maize_pare).

### RNA extraction and gene expression analysis

Total RNA was extracted from 100 mg of pooled leaf tissue from at least 3 plants using Quick-zol reagent (Kalium Tech) according to manufacturer’s instructions. Then, 1 μg of total RNA was incubated with RNase-free DNase I (1 unit/μl) following the protocol provided by the manufacturer (Promega), followed by reverse-transcription into first-strand cDNA using ImProm II reverse transcriptase (Promega), anchored oligo(dN-T_20_) and stem-loop oligonucleotides for miRNA394 when necessary (Varkonyi-Gasic et al., 2007). The obtained cDNA was used as a template for semiquantitative RT-PCR or Real Time PCR amplification in an AriaMx 1.6 (Agilent). When expression was analyzed using Real Time PCR, TransStart Green qPCR SuperMix (2X) was used, including ROX dye as internal reference and following the manufacturer’s instructions (Transgene Biotech). Primers used are reported in Table S1. Amplifications were performed under the following conditions, according to manufacturer’s manual: 30 s at 94 °C followed by 40 cycles at 94 °C for 10 s, 53 °C for 5 and 72 °C for 10 s, and a final elongation at 72°C for 20 s. Three biological replicates and three technical replicates were performed for each sample. Melting curves for each PCR product were determined by measuring the decrease of fluorescence with increasing temperature in a three-step cycle (95-65-95 °C). Analysis of the melting curve together with visualization of PCR products on a 2% (w/v) agarose gel was used to assess amplification specificity and confirm the size of the amplification products. Gene expression levels were normalized to that of maize *UBIQUITIN 1* gene (Zm*UBI1,* Zm00001d015327). Analysis was performed using the ΔCt method, after assessment of amplification efficiency close to 100 % (Livak and Schmittgen, 2001). When expression was assessed using semiquantitavie RT-PCR, PCR reactions were set up using EasyTaq (Transgene Biotech) in a mix containing 1X Taq polymerase buffer; 1.5 mM MgCl_2_; 200 μM dNTPs; 0.5 μM forward and reverse primers; 1U Taq polymerase and 1 μl cDNA template. The PCR program used was 94 °C for 2 min. followed by 28 cycles of 94 °C for 20 s; annealing temperature for 20 s (according to sequences in Table S1) and 72 °C for 30 s, and a final elongation at 72 °C for 2 min. PCR products were run in 2 % w/v agarose gels stained with Gel Red (Biotium).

### Growing conditions and drought assay

Maize plants were planted in soil using 1 l pots and transferred to a growth chamber under long-day conditions (16 h light, 28 °C/8 h dark, 26-28 °C). For control conditions, plants were watered to full field capacity every two days throughout the experiment. For drought conditions, irrigation was withheld during 25 days. Watering stopped when plants reached the V3 stage, which occurred 10 days after sowing, pots were weighted every day until a constant weight was measured for two consecutive days, which occurred 20 days after sowing, irrigation was resumed after a period of drought of 25 days (35 days after sowing). Watering was resumed during 7 days and plant survival was recorded each day to evaluate plant survival.

A similar experimental design was used for Arabidopsis plants. Arabidopsis seeds were planted in soil, stratified for 4 days at 4 °C and transferred to a growth chamber under long-day conditions (16 h light/8 h dark) at 22-24 °C. For control conditions, plants were watered regularly every two days throughout the experiment. For drought conditions, watering stopped after 15 days of plant growth in regular watering conditions, irrigation was resumed after a period of 15 days of drought and plant survival was recorded during 7 days of re-watering.

### Growth and physiological measurements

For maize plants, net photosynthesis (Pn, μmol CO_2_.m^-2^.s^-1^), leaf transpiration (E, mmol H_2_O.m^-2^.s^-1^) and stomatal conductance (gs, mmol H_2_O.m^-2^.s^-1^) were simultaneously measured with a portable infrared gas analyzer (CIRAS 2, PP Systems, Amesbury, MA, USA). The water use efficiency (WUE = Pn/gs x 1000, μmol CO_2_.μmol H_2_O^-1^) was also calculated. Plant height was measured for all plants from the base of the stem to the longest fully extended leaf. Greenness index was assessed using SPAD (Minolta), an optical device that measures light absorption of the leaf at 650 and 940 nm. The quotient of these differences is an indicator of chlorophyll content and is reported in non-dimensional SPAD units.

These traits were measured for fully irrigated control and drought-treated plants for the four analyzed genotypes 15 days after the last day of irrigation of treated plants (25 days after sowing). In all cases, equivalent fully expanded leaves were sampled and measurements were conducted on at least five plants per condition and genotype.

### Root growth and architecture

Maize roots were analyzed using WinRhizo Scanner and Root Image Analysis System (Reagent Instruments). Roots were washed and placed in a transparent tray and scanned to measure total root length, number of tips as an indication of secondary root development and total lateral root length corresponding to roots with diameters between 0 and 1 mm.

Arabidopsis root length and number of lateral roots was determined by sowing seeds in plates containing 0.5X Murashige & Skoog media (PhytoTechnology Laboratories), 8% w/V agar and supplemented with kanamycin. Plates were grown in vertical position for 15 days under growing conditions specified above.

### Chlorophyl fluorescence measurements

All spectroscopic measurements were made on equivalent fully expanded leaves from watered and drought-treated plants, on the first day of water deprivation and at 3 additional time points in a 25-days-long drought experiment. Data was collected using MultispeQ (Kuhlgert *et al*., 2016) linked to the PhotosynQ platform (http://www.photosynq.org). Measurements on maize plants were performed at 200 μmol photons m^−2^.s^−1^ red actinic light. Saturating pulses (0.5 s) of 6000 μmol photons m^−2^.s^−1^ were used to obtain maximum fluorescence signals on light-adapted states (Fm’). These measurements were used to estimate the Fv’/Fm’ parameter ((Fm’–F0′)/Fm’) which reflects the maximum efficiency of PSII in light-adapted conditions, the quantum yield of PSII (PhiII), the fraction of light dissipated as non-photochemical quenching (PhiNPQ), the total electron flow from antennae complexes into PSII (linear electron flow, LEF), the fraction of light lost through non-regulated photosynthesis inhibitor processes (PhiNO) and the fraction of open centers at PSII (qL). These photosynthetic parameters were calculated according to Baker (2008). The NPQt parameter was determined in light-adapted leaves according to Tietz *et al*. (2017).

### Determination of relative water content

Pieces of leaf blades corresponding to plants from each group (genotype x treatment; n ≥ 4 for each) were collected and immediately weighted to obtain the fresh weight (FW). The leaves were then soaked in water for 2 h, blotted with tissue paper to remove excessive moisture, and weighed to obtain the turgid weight (TW). The turgid leaves were oven-dried at 80 °C for 72 h and weighed to obtain the dry weight (DW). The relative water content (RWC) was calculated according to the following formula: RWC (%) = [(FW-DW) / (TW-DW)] ×100, (Slavik, 1974).

### Ion leakage determination

Pieces of 10 cm^2^ of leaf blades corresponding to plants from each group (genotype x treatment; n ≥ 4 for each) were collected and incubated in water at room temperature for 30 minutes with agitation, followed by measurement of initial conductivity (C_i_) using a digital conductivity meter. After autoclaving at 121 °C for 20 minutes, samples were allowed to cool down and final conductivity (C_f_) was measured. Percentual ion leakage was calculated as 100xC_i_/C_f_ for control and drought-treated samples.

### Determination of wax content

Epicuticular wax extraction was adapted from Bewick et al. (1993). For each group of plants (genotype x treatment), 15 cm measured from the distal part of the leaves were collected from each of five plants, from well-watered and drought-treated plants. Leaves were dipped (excluding the cut area) for 20 s in 50 ml chloroform contained in pre-weighted Falcon tubes (inital weight) and total submerged leaf area was calculated using ImageJ software on photographed leaves. The wax-containing chloroform solution was kept in a safety hood until complete chloroform evaporation, tubes were weighed (final weight) and wax content was determined by weight difference, and wax content per unit area was calculated.

### Statistical analysis

Two-way analysis of variance (ANOVA), using genotype and treatment as factors, were carried out to analyze interactions and main effect on the studied traits in maize mutants during drought experiments. Tukey’s post-hoc tests were used to determine the differences. For survival curves, the Kaplan-Meier log-rank analysis was used, followed by Holm-Sidak method for pairwise comparisons. Statistical evaluations were performed using R Studio and for graphical representation of the data Microsoft Excel was used.

## Acknowledgements and fundings

Authors would like to thank Marcos Reyes and Mauro Alisio, CPA members of the Argentinean Consejo Nacional de Investigaciones Científicas y Técnicas (CONICET) for collaboration with plant care and maintenance. M. D is a member of CONICET and UNL professor.

This research was supported by Fondo para la Investigación Científica y Tecnológica (FONCYT) through grants PICT 2015-0198, PICT 2018-1090 and PICT 2021-0015 to M. D.

## Authors contributions

F. M. performed all main experiments, maize plants growth, maintenance and characterization of mutant populations; A. L. conducted chlorophyl fluorescence experiments and related analyses; M.A.P. characterized transgenic Arabidopsis plants; C. B. contributed to physiological characterization of maize plants; B. M and M. C. T. provided and help characterize *zmlcr* mutants; M. D. was responsible for study conception and design, coordination of activities, manuscript writing and revision with co-authors. All authors read and approved the final manuscript.

---

**Figure S1:**
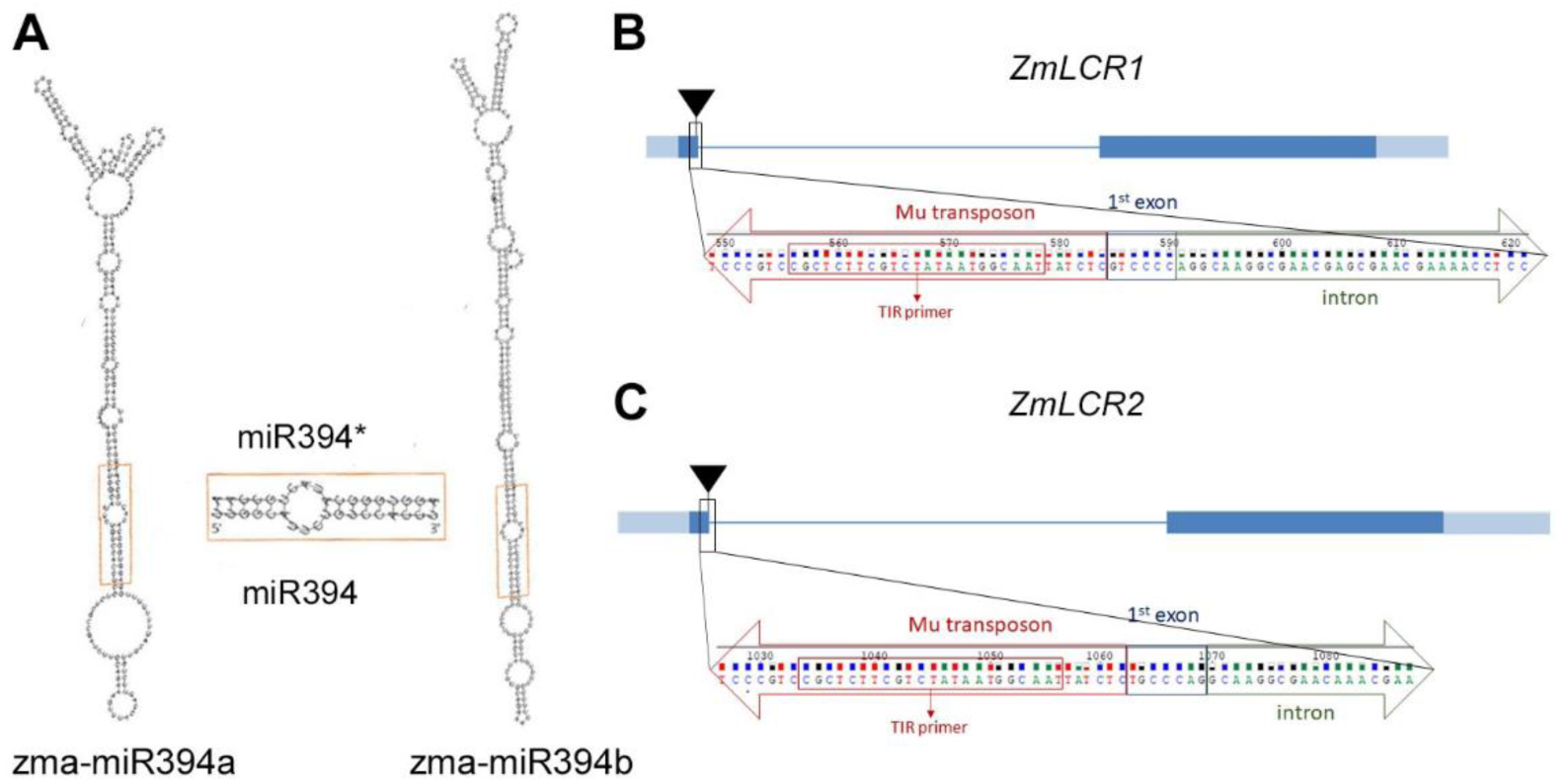
A) Hairpin structures of zma-miR394a and zma-miR394b precursors precited using RNA-fold. Duplex produced from both structures is identical as well as mature miR394 sequence. B) Representation of transposon Mu insertion in the first exon of ZmLCR1 (TUSC collection). Sequencing indicates the insertion interrupts the coding sequence of this gene. C) Representation of transposon Mu insertion in the first exon of ZmLCR2 (UniforMu collection). Sequencing indicates the insertion interrupts the coding sequence of this gene.

**Figure S2:**
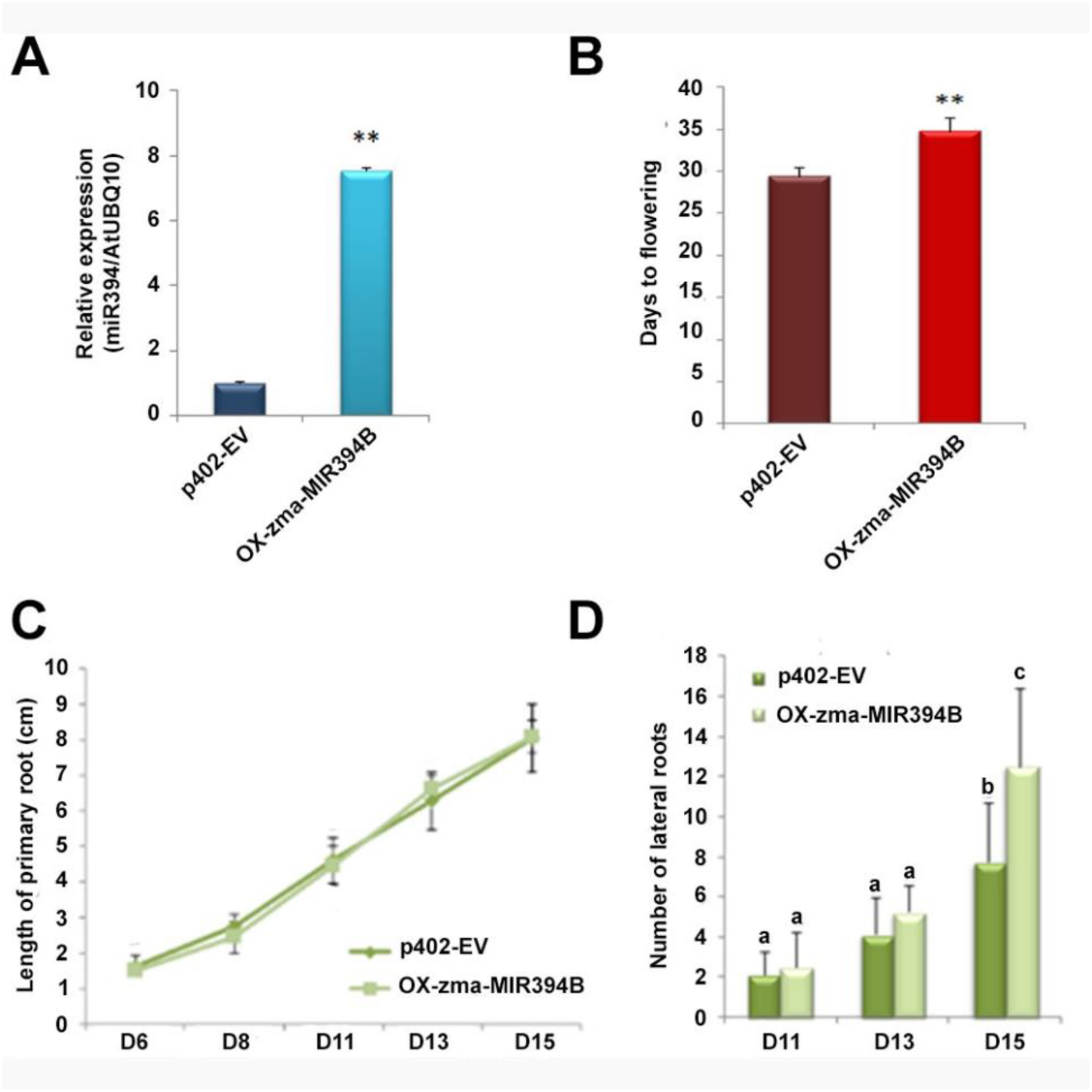
Characterization of Arabidopsis *OX-zma-MIR394B* plants. A) Relative expression of mature miR394 in control p402-EV plants transformed with empty pGWB402Ω vector and *OX-zma-MIR394B* plants overexpressing zma-MIR394B gene. (AtUBQ10 gene expression was used to normalize miR394 expression; mean ± SD is represented; ** = p < 0.001 according to Student’s t-test). B) Flowering time in p402-EV and *OX-zma-MIR394B* plants (mean ± SD is represented; ** = p < 0.001 according to Student’s t-test; n ≥ 10). C) Length of primary roots in p402-EV and *OX-zma-MIR394B* plants grown *in vitro* in vertical plates, determined at 6, 8, 11, 13 and 15 days of growth. No significant differences were observed between genotypes (ANOVA, P < 0.05). D) Number of lateral roots in p402-EV and *OX-zma-MIR394B* plants grown *in vitro* in vertical plates, determined at 11, 13 and 15 days of growth. Different letters indicate significant differences (Two-way ANOVA followed by Tukeýs post hoc test, P < 0.05).

**Figure S3:**
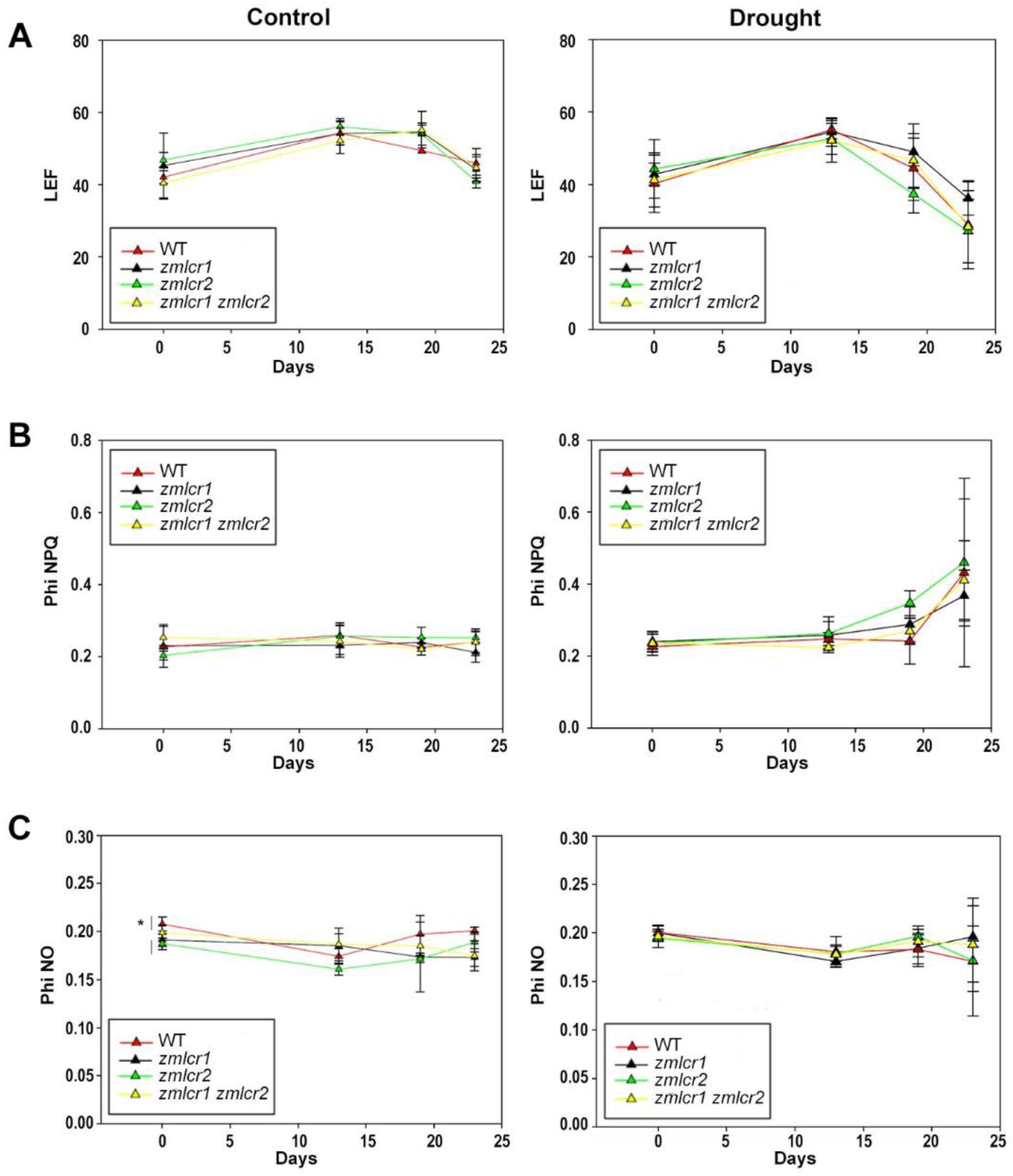
Energy management at PSII in WT, *zmlcr1, zmlcr2* and *zmlcr1 zmlcr2* mutants. Evolution of LEF (Linear Electron Flow), PhiNPQ and PhiNO in fully-irrigated plants (Control) and water-deprived (Drought).

**Table S1:** Primers used.

**Table S2:** Genomic coordinates of the miR394 pathway components in maize.

|  | Gene ID<br>B73 RefGen v3 | Gene ID<br>B73 RefGen v5 | Genome location<br>B73 RefGen v5 |
| --- | --- | --- | --- |
| *zma-MIR394a* | GRMZM5G817717 | Not annotated | Chr5:198202923..198203048 |
| *zma-MIR394b* | GRMZM5G871952 | Not annotated | Chr4:159533569..159533690 |
| *ZmLCR1* | GRMZM2G119650 | Zm00001eb143020 | Chr3:160877675..160881388 |
| *ZmLCR2* | GRMZM2G064954 | Zm00001eb364210 | Chr8:167387233..167391418 |

**Table S3:** Fv’/Fm’.

|  | WT | *zmlcr1* | *zmlcr2* | *zmlcr1 zmlcr2* |
| --- | --- | --- | --- | --- |
| **D0 - Control** | 0.693 ± 0.017 | 0.689 ± 0.028 | 0.700 ± 0.004 | 0.682 ± 0.011 |
| **D0 - Drought** | 0.696 ± 0.004 | 0.689 ± 0.009 | 0.689 ± 0.014 | 0.689 ± 0.010 |
| **D23 - Control** | 0.689 ± 0.014 | 0.686 ± 0.019 | 0.684 ± 0.015 | 0.671 ± 0.029 |
| **D23 - Drought** | 0.564 ± 0.158 | 0.610 ± 0.071 | 0.507 ± 0.107 | 0.590 ± 0.074 |

